# Characterization of Prenylated C-terminal Peptides Using a Novel Capture Technique Coupled with LCMS

**DOI:** 10.1101/2020.01.15.908152

**Authors:** James A. Wilkins, Krista Kaasik, Robert J. Chalkley, Al L. Burlingame

## Abstract

Post-translational modifications play a critical and diverse role in regulating cellular activities. Despite their fundamentally important role in cellular function, there has been no report to date of an effective generalized approach to the targeting, extraction and characterization of the critical c-terminal regions of natively prenylated proteins. Various chemical modification and metabolic labelling strategies in cell culture have been reported. However, their applicability is limited to cell culture systems and does not allow for analysis of tissue samples. The chemical characteristics (hydrophobicity, low abundance, highly basic charge) of many of the c-terminal regions of prenylated proteins have impaired the use of standard proteomic workflows. In this context, we sought a direct approach to the problem in order to examine these proteins in tissue without the use of labelling. Here we demonstrate that prenylated proteins can be captured on chromatographic resins functionalized with mixed disulfide functions. Protease treatment of resin-bound proteins using chymotryptic digestion revealed peptides from many known prenylated proteins. Exposure of the protease-treated resin to reducing agents and hydro organic mixtures released c-terminal peptides with intact prenyl groups along with other enzymatic modifications expected in this protein family. Database and search parameters were selected to allow for c-terminal modifications unique to these molecules such as CAAX box processing and c-terminal methylation. In summary, we present a direct approach to enrich and obtain information at a molecular level of detail about prenylation of proteins from tissue and cell extracts using high performance LCMS without the need for metabolic labeling and derivatization.

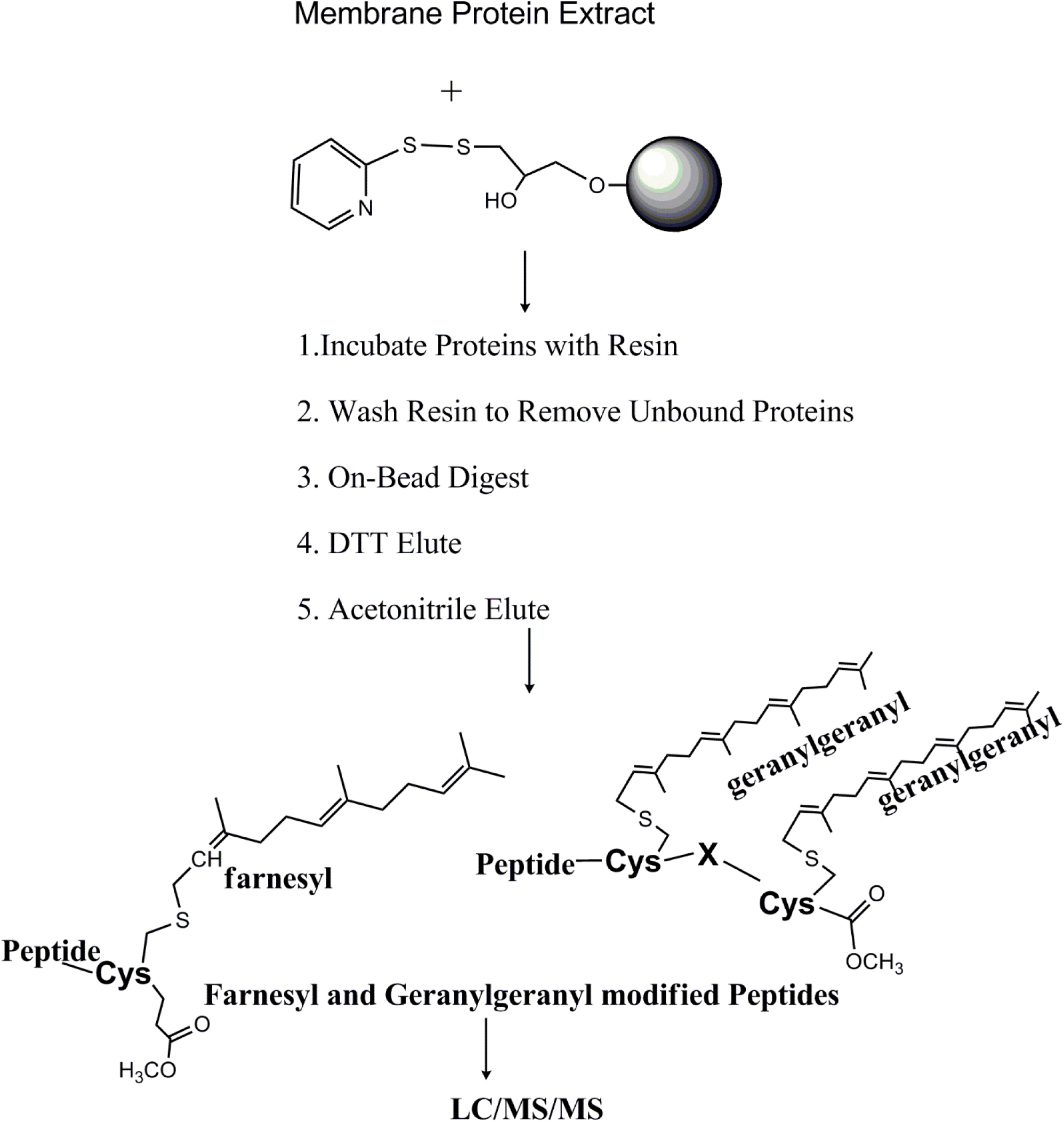

## INTRODUCTION

An ever-increasing array of proteomics methods using targeted qualitative and quantitative approaches aimed at characterizing and quantifying posttranslational modifications have become feasible using high performance LCMS methodologies and current instrumentation. This has enabled the detection, characterization and quantitation of proteins and their modifications ranging from phosphorylation to glycosylation among many others on a global, proteome-wide scale. For more comprehensive results methods often leverage selective enrichment of a targeted sub-stoichiometric modification using antibody or other affinity-based recognition techniques as well as a variety of chemical targeting techniques. Such enrichment mixtures are separated and analyzed by reversed phase chromatography and high sensitivity mass spectrometry (1, 2). A unique group of membrane-associated intracellular proteins are post-translationally modified at cysteine residues near their c-termini with either C15 (farnesyl) or C20 (geranylgeranyl) isoprenoid moieties via a thioether linkage. “Prenylation” of target proteins is necessary for their association with the plasma membrane and with membranes of cellular organelles, and thus participates in specifying their cellular localization and function. Many of them are GTPase switches and play a central role in cell signaling, transport, and cytoskeletal rearrangement among other activities. Examples include the RAS family: KRAS4A and 4B (human), Hras and Nras. RAS proteins trigger cell growth through kinase cascades. Posttranslational modifications of these proteins including the enzymes involved in prenylation and other c-terminal processing have been described (3–5) Point mutations in the GTP/GDP binding region of RAS proteins are known to play a causative role in a variety of human cancers; mutations may also be responsible for other metabolic problems associated with autism and other neurological disorders (6, 7).

Prenylation of protein targets occurs via enzymatic conversions catalyzed by one of three enzymes. Farnesyltransferase (FTase) and geranylgeranyl transferase I (GGTase I) are responsible for all reactions involving proteins with a c-terminal “CAAX” motif. GGTase II catalyzes transfer of the C20 polyisoprene to substrate Rab and other proteins with various c-terminal cysteine motifs including XXCC, XXCXC, XXCCX, XCCXX, XCXXX and CCXXX where “X” is any amino acid (8, 9). Several accessory proteins are known to play roles in the interaction of GGTase II with its substrates (10). In addition to the preceding, some proteins, such as those belonging to the Ras family (e.g. Kras, Nras and Hras) are predicted to contain palmitoyl or other fatty acyl groups attached through cysteine thioesters located in their c-terminal regions. The enzymatically-catalyzed addition of farnesene (C15) or geranylgeranyl (C20) to cysteine residues located in the “CAAX” motif at the C-terminus of target proteins is absolutely required for their normal cellular function and distribution at the plasma membrane and other membrane locations. Other mandatory enzymatic processing of these so-called “CAAX box” proteins includes enzymatic removal of the c-terminal “AAX” sequence catalyzed by Ras converting enzyme I (Rce I) and c-terminal methyl esterification of the resulting mature protein by isoprenylcysteine carboxymethyltransferase (Icmt).

Prenylated proteins have been shown to be excluded from lipid raft structures in the plasma membrane (11) but some proteins such as Hras carry both prenyl and palmitoyl modifications, perhaps rendering their membrane associations more complex (12). Modifications of the c-terminal amino acid sequence of KRAS 4B were recently shown to exert a profound effect on its membrane interactions (13). Targeted farnesyl transferase inhibitors have been tested in a clinical setting against KRAS mediated cancers. However, it was discovered that inhibition led to a cellular switch to geranylgeranylation of the protein, presumably by GGTase 1 (14), rendering the drugs ineffective in this setting, although other clinical targets such as the laminopathies (15) including Hutchinson-Gilford progeria syndrome have proven more promising in this regard.

Direct characterization of prenylation has been limited by the lack of suitable analytical approaches, although various labeling and capture strategies in tissue culture have been used (16), usually employing isotopically labelled isoprenoid precursors or covalent handles linked to precursors in cell culture systems (17, 18). Heretofore, direct isolation and characterization of prenyl proteins and their c-terminal peptides has proved to be particularly difficult due to several factors: 1. c-terminal peptides are unusually hydrophobic rendering them poor candidates for standard (C18) reversed phase separations. 2. Many of these proteins contain a large number of lysine and arginine residues near their c-terminus that are known to promote interaction with the inner leaflet of the plasma membrane (4) but, more importantly in this context, discourage the use of tryptic protein digestion, since resulting peptides would be too short (and predicted to be singly charged) eliminating meaningful analysis by LCMSMS in many cases. As it is, many of these peptides such as the c-terminal (human) KRAS 4B chymotryptic peptide with 14 basic residues in a total of 25 amino acids present a unique analytical challenge. 3. Customized informatics (as described below) is required in order to effectively detect the presence of these peptides in database searches. 4. Up to now, there were no “targeted” or chemistry-based affinity methods for peptides containing the isoprene moiety. A recent report described an elegant labelling strategy with chemically reactive isoprenoid probes designed to produce reporter ions in mass spectrometry in which a number of prenyl proteins and their c-terminal peptides were identified and characterized (19). However, to characterize these proteins and their various posttranslational modifications from tissue and particularly to enable protein analysis from normal and disease states, a method that can enrich natively modified protein is necessary.

We have approached this problem using selective membrane protein extraction coupled with a new method that enables targeted capture of prenylated proteins.

## EXPERIMENTAL PROCEDURES

### Experimental Design and Statistical Rationale

This study details results obtained from 4 different mouse brain extracts, thus 4 biological replicates. Within each replicate, we analyzed total proteins captured (chymotryptic digest) as well as peptides eluted either by exposing the capture matrix to reducing conditions, nonpolar solvent or a combination of the two.

## Methods

### Extraction and Processing of Prenylated Proteins from Mouse Brain Tissue

#### Mouse brain sample preparation

Mice were housed a 12-h light-dark cycle and all studies were approved by the Institutional Animal Care and Use Committee at UCSF. Mouse brain tissue was dissected from adult mice and kept frozen in dry ice or liquid nitrogen. In each experiment washed membranes were prepared from 3 mouse brains using a hypotonic, acidic buffer (0.01N HCl) to neutralize endogenous serine protease activity with mechanical disruption using a Dounce homogenizer as outlined in figure S1. The homogenate (50ml) was centrifuged at 10,000xg for 10 minutes at 4°C. The supernatant was discarded, membranes were resuspended in the same volume of 0.01N HCl and the suspension centrifuged as above. After another wash with 0.01N HCl, membranes were extracted with 5ml of RIPA extraction buffer (50mM Tris base, 150mM NaCl, 1% (w/v) Igepal-630, 12mM sodium deoxycholate, 0.1% (w/v) sodium dodecyl sulfate, pH 7.6) for 1.5 hours at 4°C with end-over-end mixing. Extracted membranes were sedimented at 10,000xg for 10 min at 4°C and a second extraction was performed under the same conditions for 1.5 hour (figure S1). Following sedimentation of the extracted membranes as above, the 2 soluble extracts were combined, and the solution was adjusted to 10mM tris carboxyethyl phosphine (TCEP) using a 1M stock solution at 4°C. A protein-containing precipitate immediately formed, and the suspension was incubated at 4°C for 1 hour. Iodoacetamide solution in water (0.5M) was added to achieve a final concentration of 10mM and the solution was incubated at room temperature in the dark for 30 minutes. Finally, the suspension was centrifuged at 10,000xg for 10 minutes. Insoluble proteins were recovered and resuspended in 5 ml of a solution containing 8M urea and 50mM ammonium bicarbonate (ABC), pH 7.8; the supernatant (containing most of the RIPA buffer detergents) was discarded. Solubilized proteins were exchanged into the same buffer (8M urea/50mM ABC) using gel permeation chromatography on Sephadex G-25 to remove residual TCEP and iodoacetamide.

#### Protein capture and targeted peptide strategy

Prenyl proteins were captured on thiopropyl Sepharose 6B as follows. Thirty-five mg of lyophilized resin was suspended in 1ml of water and the suspension was gently mixed for 10 minutes. The swollen resin was then washed in a spin column using 5 rinses of 0.5 ml water followed by 5 rinses of 0.5ml of 50mM ABC. For each wash, the resin was thoroughly resuspended before centrifugation. The resin was then mixed with 1mg of the urea-containing protein solution from the G25 column and the mixture was incubated on an end-over-end mixer for 18 hr at 4 degrees C. Unbound proteins were removed from the thiopropyl resin using a spin column followed by extensive washing using 0.5ml volumes (x5) of the following: 6M urea/50mM ABC; 2m NaCl; 70% acetonitrile/ 0.1% formic acid in water; water, 50mM ABC. The washed resin was then resuspended in 200 microliters of 50mM ABC. One microgram of chymotrypsin (Promega sequencing grade) was added to the suspension and the mixture was incubated with vigorous mixing at 37°C for 5-6 hours. The resin was placed in the spin column and unbound chymotryptic peptides were isolated. The resin was washed with 100 microliters of 0.1% formic acid and the wash was combined with the released peptides. Unbound chymotryptic peptides were saved for analysis.

#### Peptide elution with dithiothreitol (DTT) followed by 50% acetonitrile

Prenylated peptides were released from the resin using a 2-step method. All separations were done using spin columns. In the first step (DTT elution), resin was incubated with 200 microliters 50mM ABC containing 10mM DTT for 2 hours at room temperature with mixing. This step cleaved the thiopyridine protecting groups from the resin and released some prenylated peptides. Following incubation, the resin was separated from the DTT-eluted solution followed by a rinse with 100 microliters of 0.1% formic acid in water (which was combined with the DTT eluted peptides). The resin was then suspended in 200 microliters of 50% acetonitrile/50% water containing 0.1% formic acid and incubated as above (50% ACN elution) and the suspension was mixed overnight at room temperature. The resin was rinsed with 100ul of the same solution and the rinse was combined with the eluted peptides from this step.

Peptides from the DTT and 50% ACN elutions were dried under vacuum and dry material from each fraction was resuspended in 50 microliters of 1 % formic acid in water. Peptides were purified by removing salt and reagents using C18 containing pipet tips (OMIX, Agilent) and eluted from the tips using 50% acetonitrile, 0.1% formic acid in water. Eluted peptides were dried under vacuum and resuspended in 10% acetonitrile, 0.1% formic acid in water for analysis by LCMS.

#### Liquid Chromatography/Mass Spectrometry Analysis

HCD data on samples was obtained using a Q Exactive Plus mass spectrometer (Thermo Scientific) equipped with a nano-Acquity UPLC system (Waters, Milford, MA). Peptides were fractionated on a 20 cm × 75 μm ID Picofrit (New Objective) column packed with 3.5 micron, 100 Å pore size Kromasil C18 using linear gradients of solvents (defined in the figure legends) composed by mixing solvent A: water + 0.1% formic acid and solvent B: acetonitrile + 0.1% formic acid. EThcD data were acquired using a Fusion Lumos (Thermo Scientific) mass spectrometer. In this case, peptides were fractionated on a 15cm Easyspray (Thermo Scientific), C18 column. All mass measurements were performed in the Orbitrap. The 10 most abundant multiply charged ions were computer-selected for HCD analysis in the Q Exactive; the Fusion Lumos was programmed to select the top 20 most abundant ions. The trigger intensity was set to 2000. HCD and EThcD fragments were measured in the Orbitrap.

#### Prenyl Protein Database

In order to create a database of possible prenylated proteins we first searched the *Mus musculus* protein database (UniProtKB.2015.12.1) using the known sequence motifs as templates using MS-Pattern in Protein Prospector (ref PMID:16401513). We then filtered this list to only those annotated as being prenylated according to Uniprot), to give a total of 284 entries.

#### Database Searching

Peaklists were extracted using an in-house program and data was searched against the Uniprot *Mus musculus* database (73,955 entries, downloaded) (and concatenated with a randomized sequence for each entry) or against an in-house produced accession list (see below) using Protein Prospector (version 6.1.22). To allow for cleavage of the “AAX” motif at the C-terminal side of cysteine by RceI in addition to chymotryptic cleavage the enzyme specificity searched was cleavage C-terminal to Phe, Tyr, Trp, Leu or Cys, allowing for 3 missed cleavages and one end non-specific because of the loose specificity of chymotrypsin. Farnesyl, geranylgeranyl and their neutral losses, carbamidomethylation of cysteine, acetylation of protein N termini, oxidation of Met, cyclization of N-terminal Gln and C-terminal methylation were allowed as variable modifications, with three modifications per peptide permitted. The required mass accuracy was 20 ppm for precursor ions, and 30 ppm for HCD and EThcD fragments. Spectra identified as representing peptides featuring a Cys-modification with a maximum E value of 0.05 and with a SLIP (site localization) score (ref PMID:21490164) of 5 were used as a starting point for further analysis. All prenylated peptide spectra were manually examined for quality and authenticity.

## RESULTS

Isoprenylated proteins associate with cellular membranes. We started our isolation procedure by preparing a crude membrane fraction from mouse brain tissue. Membranes were washed extensively to remove soluble proteins and then extracted to solubilize membrane proteins including prenylated species (fig. S1). Initially, tryptic and chymotryptic digests of the solubilized proteins were analyzed using one-dimensional LC-MS/MS. Protein Prospector was used to search peak lists using an in-house curated prenyl protein database (see methods section for construction) to speed processing. Many peptides derived from known prenylated proteins were identified in the searches (data not shown). However, like others (e.g. (20)), we were unable to detect the c-terminal peptides containing the prenyl modifications from any of the proteins identified.

While attempting derivatization methods to convert the prenyl groups into sulfhydryl-containing moieties that could be enriched using thiopropyl Sepharose (Fig 1), we discovered that the beads were able to enrich underivatized prenylated proteins from complex mixtures. After overnight incubation of the resin with protein extracts, the resin was washed extensively to remove nonspecifically bound proteins. Treatment of resin-bound proteins with chymotrypsin released many peptides including a large number from prenylated proteins, but almost no prenylated c-terminal peptides could be identified in the chymotryptic digest. However, we found that some prenylated peptides (along with other peptides from both prenyl- and non-prenyl proteins) were released from the thiopropyl resin after incubation with 10mM dithiothreitol. Elution with aqueous 50% acetonitrile containing 0.1% (v/v) formic acid released more peptides (see Table 1 and Table 2), suggesting a non-covalent hydrophobic binding mechanism, which was disrupted either by cleavage of the disulfide bond on the resin by DTT or by treatment with elevated levels of organic solvent. Most of the peptides released in these elutions were not from prenyl proteins, as revealed by searches of the entire *Mus musculus* (UniProtKB.2015.12.1) database; prenyl proteins only represented about 3-5% of the total number identified. However, this represented a significant enrichment for prenylated species compared to a proteomic survey of the crude membrane extract as stated above, and allowed detection of natively prenylated c-terminal peptides from a complex mixture for the first time. When the thiopropyl resin was pre-treated with DTT before the protein capture step, no prenylated peptides were observed in the DTT or acetonitrile eluates (data not shown), indicating the thiopyridyl leaving group is required for the binding. When we tried experiments with a related resin containing a larger, more polar glutathione linker to a thiopyridyl group this led to many fewer identifications of modified c-terminal regions. Therefore, the thiopyridyl group is not sufficient; the propyl group is also playing a role in the enrichment.

**Table 1.**
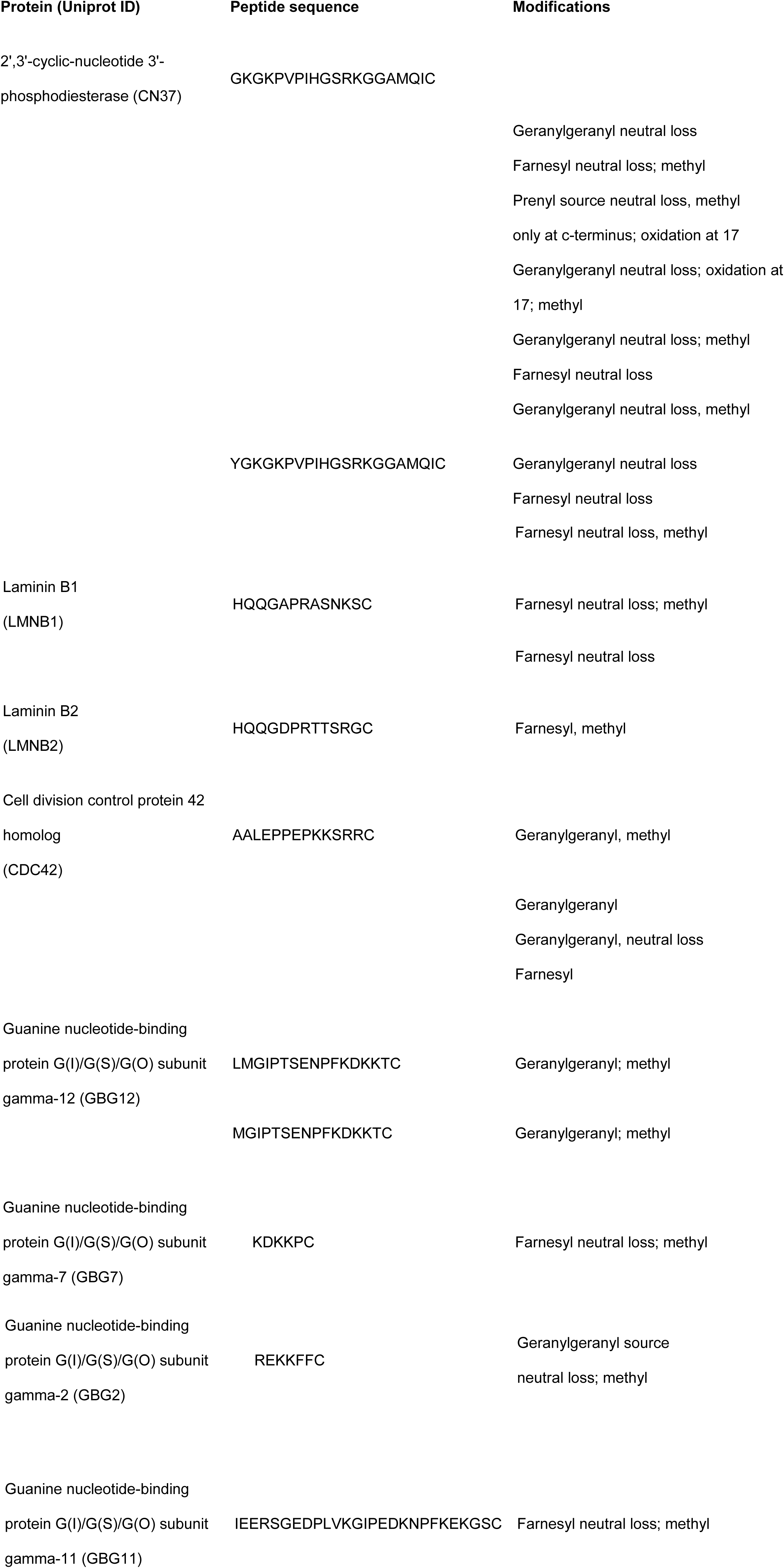
List of c-terminal modified peptides identified in a representative experiment showing multiple types of modification on some c-termini.

**Table 2.**
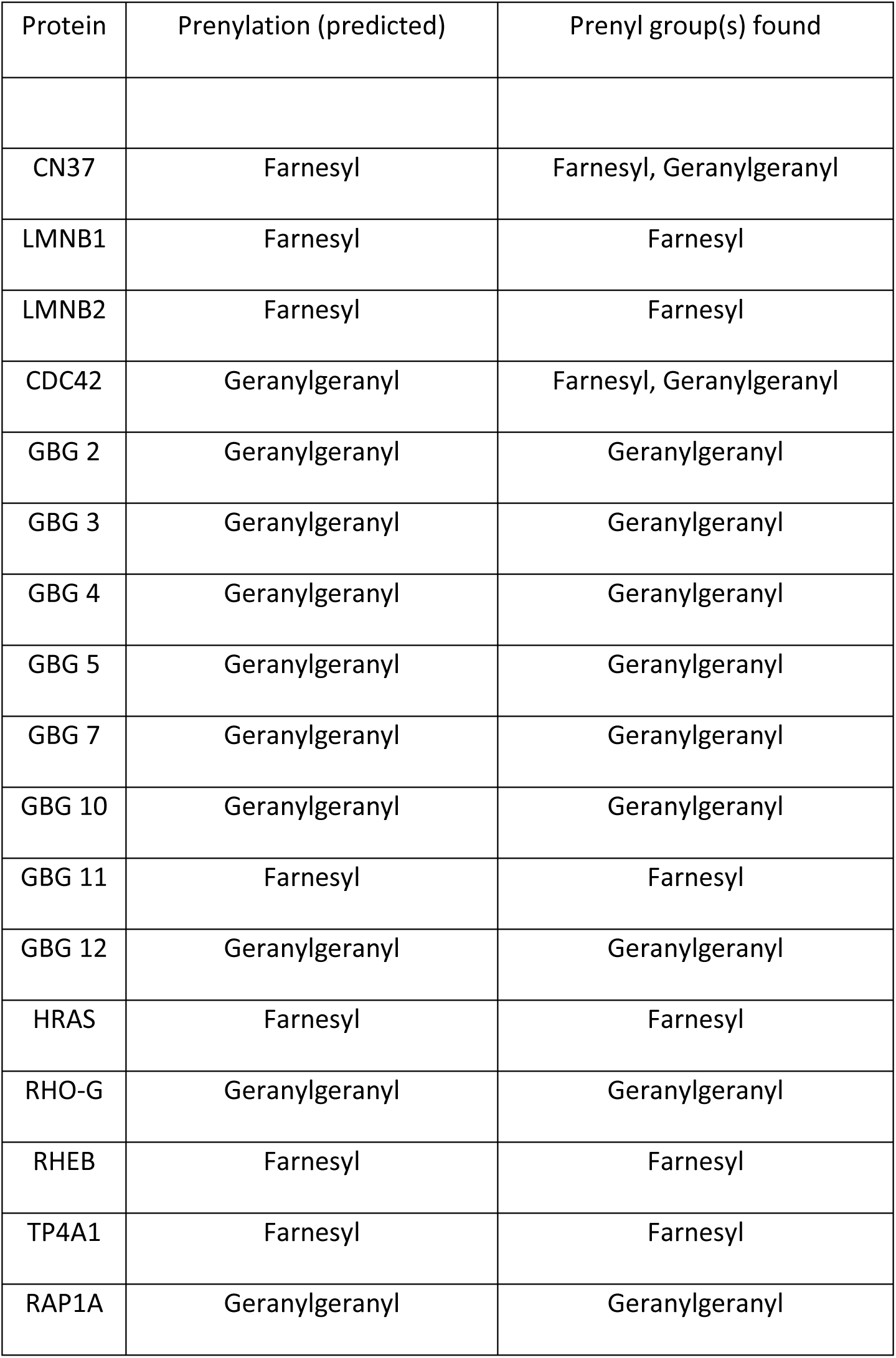
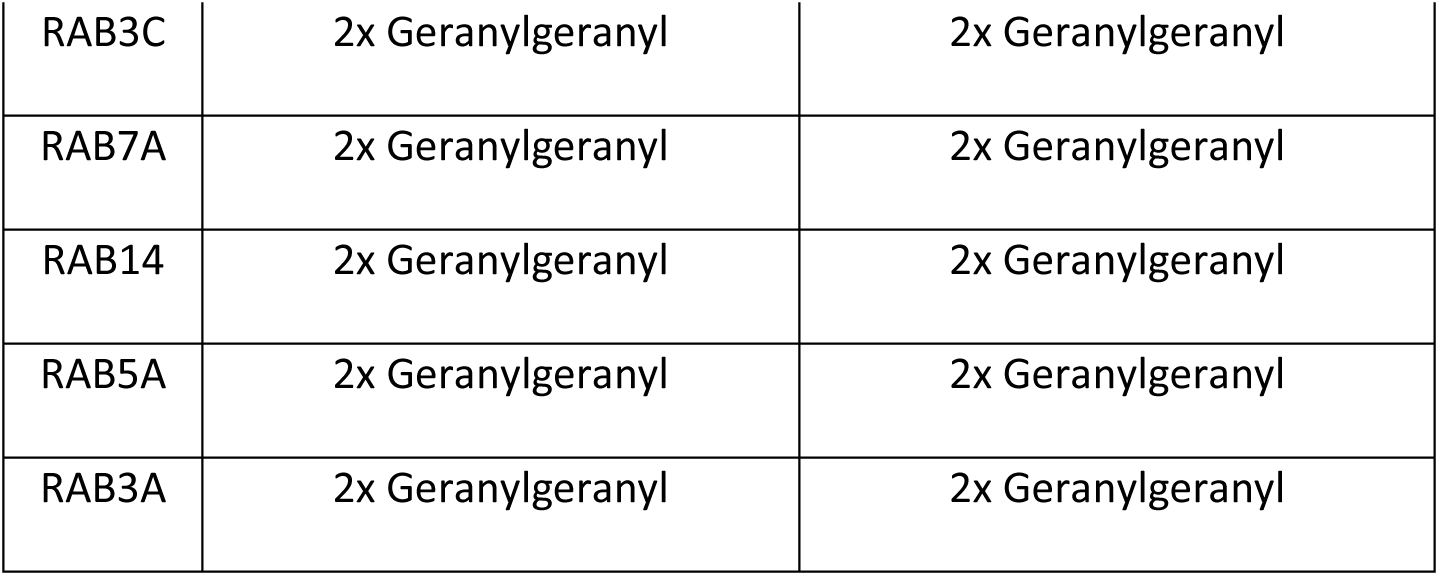
List of all c-terminal peptides identified along with their predicted (from Uniprot annotations) and found isoprene modifications

| Protein | Prenylation (predicted) | Prenyl group(s) found |
| --- | --- | --- |
| CN37 | Farnesyl | Farnesyl, Geranylgeranyl |
| LMNB1 | Farnesyl | Farnesyl |
| LMNB2 | Farnesyl | Farnesyl |
| CDC42 | Geranylgeranyl | Farnesyl, Geranylgeranyl |
| GBG 2 | Geranylgeranyl | Geranylgeranyl |
| GBG 3 | Geranylgeranyl | Geranylgeranyl |
| GBG 4 | Geranylgeranyl | Geranylgeranyl |
| GBG 5 | Geranylgeranyl | Geranylgeranyl |
| GBG 7 | Geranylgeranyl | Geranylgeranyl |
| GBG 10 | Geranylgeranyl | Geranylgeranyl |
| GBG 11 | Farnesyl | Farnesyl |
| GBG 12 | Geranylgeranyl | Geranylgeranyl |
| HRAS | Farnesyl | Farnesyl |
| RHO-G | Geranylgeranyl | Geranylgeranyl |
| RHEB | Farnesyl | Farnesyl |
| TP4A1 | Farnesyl | Farnesyl |
| RAP1A | Geranylgeranyl | Geranylgeranyl |
| RAB3C | 2x Geranylgeranyl | 2x Geranylgeranyl |
| RAB7A | 2x Geranylgeranyl | 2x Geranylgeranyl |
| RAB14 | 2x Geranylgeranyl | 2x Geranylgeranyl |
| RAB5A | 2x Geranylgeranyl | 2x Geranylgeranyl |
| RAB3A | 2x Geranylgeranyl | 2x Geranylgeranyl |

**Figure 1.**
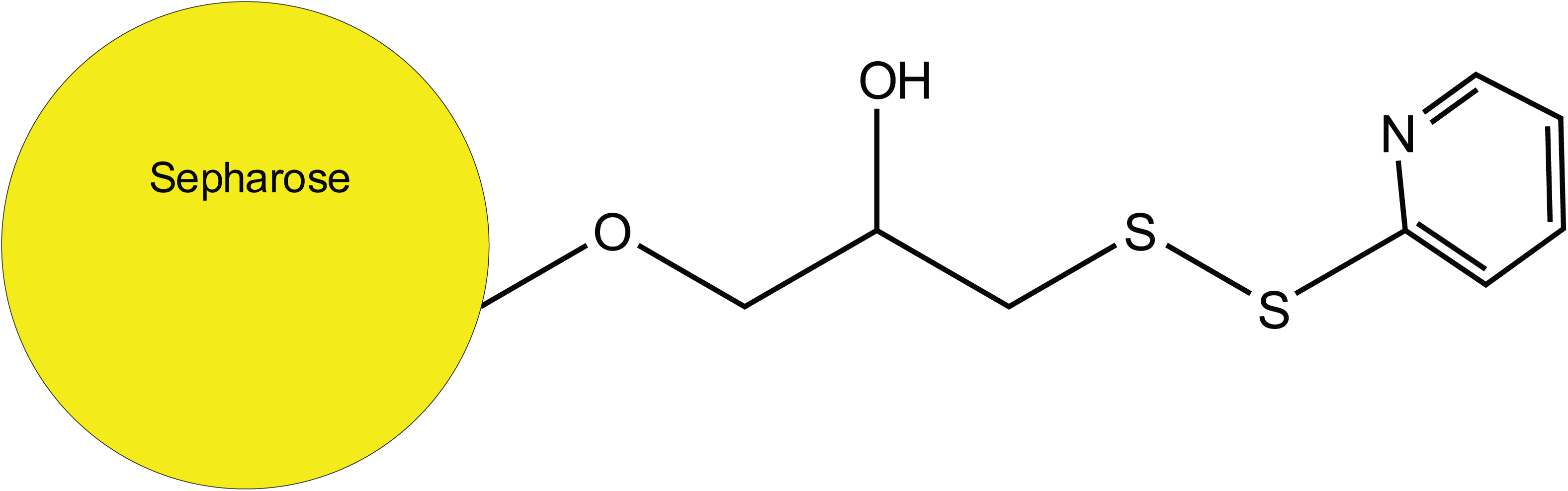
Structure of the thiopyridyl-protected thiopropyl group on thiopropyl Sepharose

All isoprene-modified peptides identified had long retention times (>60 minutes) in 2-hour liquid chromatographic separations on C18 silica-based resins, as expected given their hydrophobic character. Elution of the methylated, geranylgeranylated form of the M+4H+ peptide YGKGKPVPIHGSRKGGAMQIC (m/z 618.3604) from C18 occurred (Fig 2a) at Rt= 86.1 minutes in a 118-minute gradient from 10 to 40% acetonitrile. Figure 2b shows the M SMS spectrum of this peptide which underwent a facile loss of geranylgeranyl. The low mass region of the HCD spectrum (Figure 3) also shows characteristic fragments of the geranylgeranyl moiety at m/z 109.1, 123.12, 129.10 and 149.13, produced by fragmentation of the isoprene backbone (21). The fragments provided a convenient confirmation that spectra were of geranylgeranylated peptides and enabled detection of c-terminal peptides from Hras and from Rho G using the MS Filter utility available as part of the Protein Prospector package (see Figures S2 and S3). The c-terminal peptide from myelin-associated 2’,3’-cyclic-nucleotide 3’-phosphodiesterase (CN37) was identified farnesylated or geranylgeranylated. Table 1 presents a list of c-terminal modified peptides identified in a representative experiment showing the multiple types of modification on some c-termini. Figure 4 shows a comparison of the retention times of two versions of a CN37 c-terminal peptide (both methylated and either farnesylated (4B) or geranylgeranylated (4C)) detected in the same run. Figure 4a shows the extracted ion chromatogram of a peptide with a mass corresponding to a methyl only modification with an identical retention time to the geranylgeranylated peptide, suggesting that in this case, the methylated, geranylgeranylated peptide underwent a neutral loss of geranylgeranyl in the source of the mass spectrometer.

**Figure 2.**
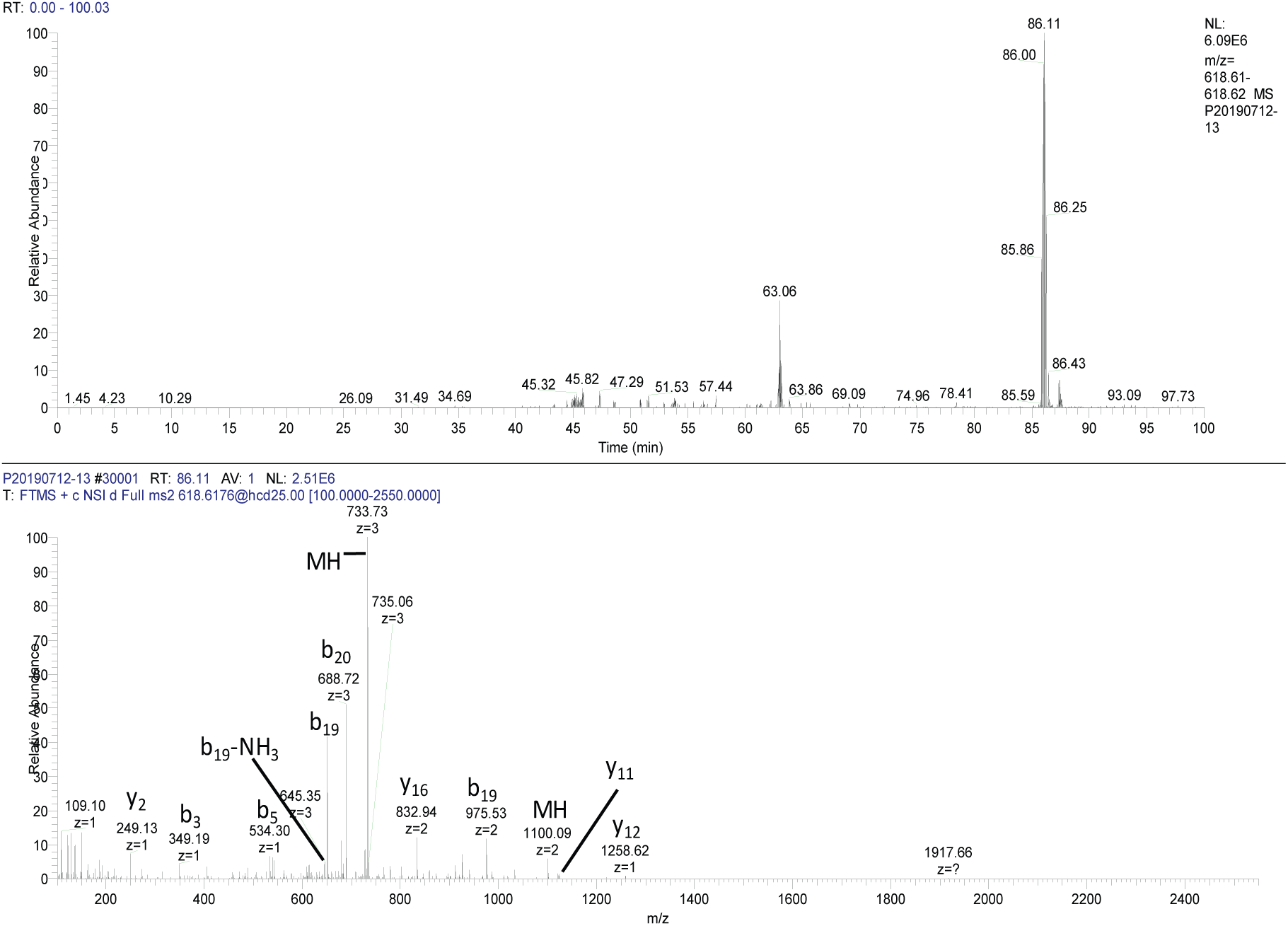
A. Extracted ion chromatogram of YGKGKPVPIHGSRKGGAMQIC-Methyl+GeranylGeranyl+4 (m/z 618.36; Rt= 86.11 min) from lc/ms separation of peptides eluted with 10mM DTT from thiopropyl Sepharose. The online reversed phase column was eluted with a linear gradient composed of water and acetonitrile each containing 0.1% formic acid from 10-40% acetoinitrile. B. ms/ms spectrum of m/z 618.36.

**Figure 3.**
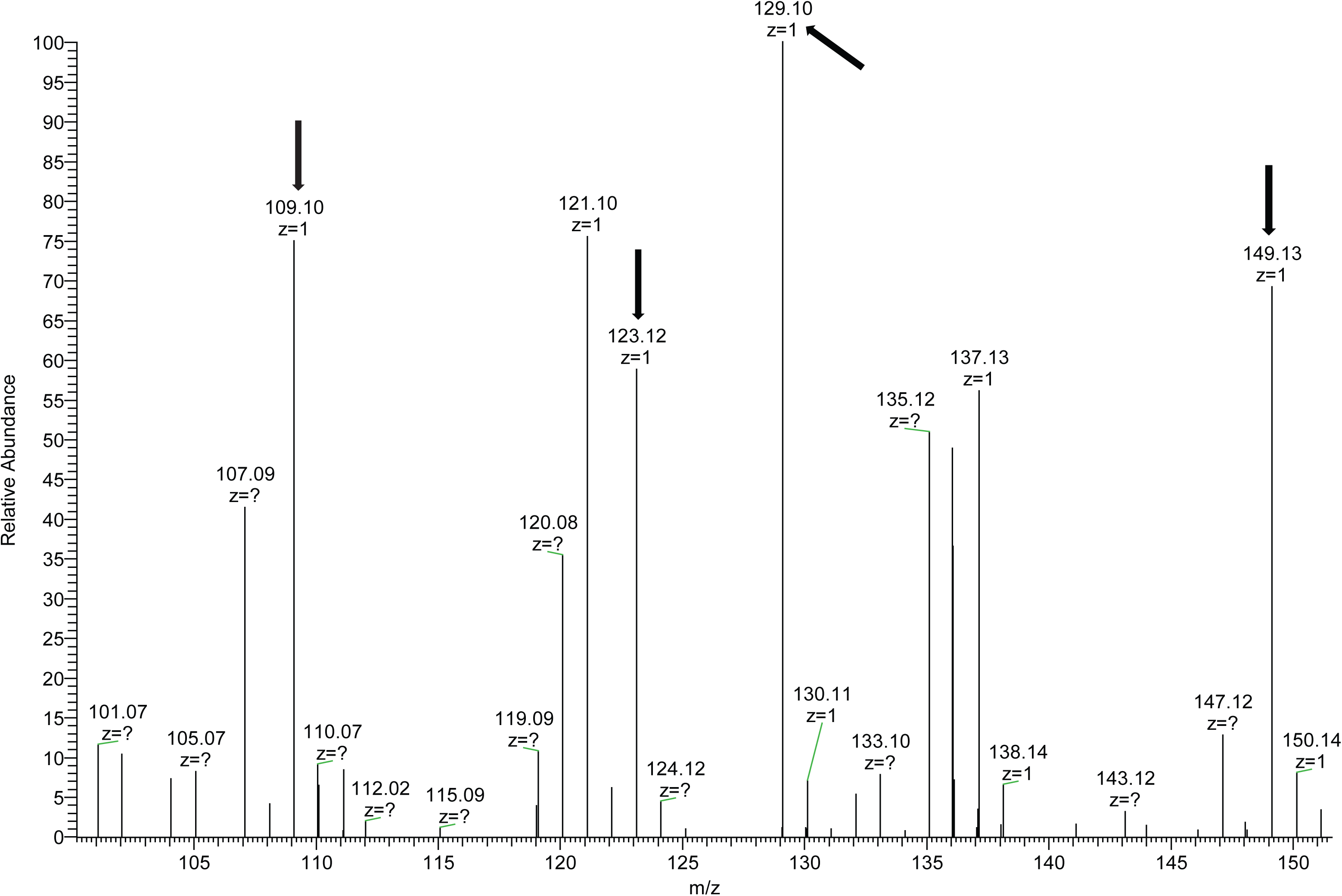
Low mass region of the msms spectrum presented in fig. 2 showing fragments of the geranylgeranyl group generated as a result of the neutral loss.

**Figure 4.**
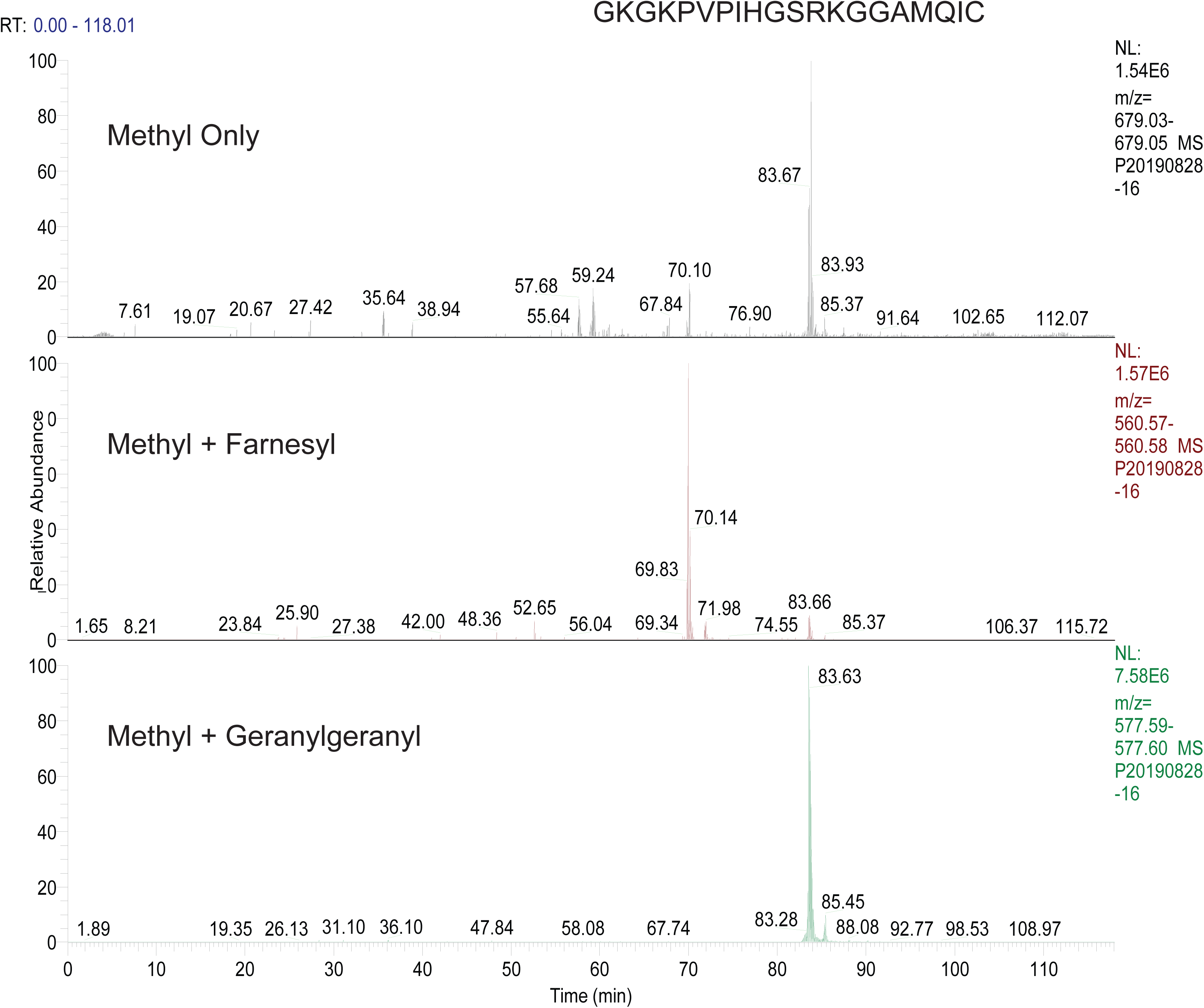
Chromatographic behavior of CN37 c-terminal peptide modified by methyl (top panel) methyl + farnesyl (middle panel) or methyl + geranylgeranyl (bottom panel) in a single run.

Prenylated c-terminal peptides were found from small guanine nucleotide binding proteins (GBG10, GBG11, GBG12, GBG7, GBG3, GBG4, GBG5) and Table 2 presents a list of all c-terminal peptides found to date along with their predicted (from Uniprot) and found prenylations. Peptides were also found from the Rab family of proteins known to be modified by the GGTase 2 enzyme (i.e. proteins containing 2 geranylgeranylated cysteines; see below). Table S1 shows a list of all prenyl proteins identified after enrichment in the four biological replicate samples, indicating a high overlap in enriched proteins.

In earlier analyses we did not identify any c-terminal peptides from doubly geranylgeranylated peptides, even though we knew through detection of other, unmodified peptides from these proteins that we were enriching proteins that had this modification. Speculating that this may be due to them being very hydrophobic we performed one more experiment. This time we analyzed the peptide mixture using a longer (155 minute) gradient running from 20% to 60% acetonitrile; i.e starting and ending at higher acetonitrile levels than usual. For this experiment, we also used a Fusion Lumos mass spectrometer equipped with EThcD to see if the different fragmentation was beneficial. Electron transfer dissociation has been shown to be particularly effective in the fragmentation of highly charged peptides, which many of the c-terminally prenylated peptides typically are ((22)). We found that the combined power of modified chromatographic conditions with the enhanced features of the Fusion Lumos instrument enabled detection of several c-termini from Rab proteins including Rab3A, Rab3C, Rab5A, Rab7A and Rab14. Figure 5 shows the elution of the 18 amino acid-doubly geranylgeranylated c-terminal peptide (GVDLTEPAQPARSQCCSN+ 3H^+^) from Rab5A. The retention time of this peptide was 138.7 min in the 155min gradient. As shown in the accompanying MSMS spectrum (panel C), fragmentation produced exclusively b and c ions and showed a neutral loss of one of the two geranylgeranyl groups. Fragments of the isoprene similar to those seen in Figure 3 were present in the low mass region of the spectrum including a peak corresponding to the intact geranylgeranyl moiety at m/z 273.25.

**Figure 5.**
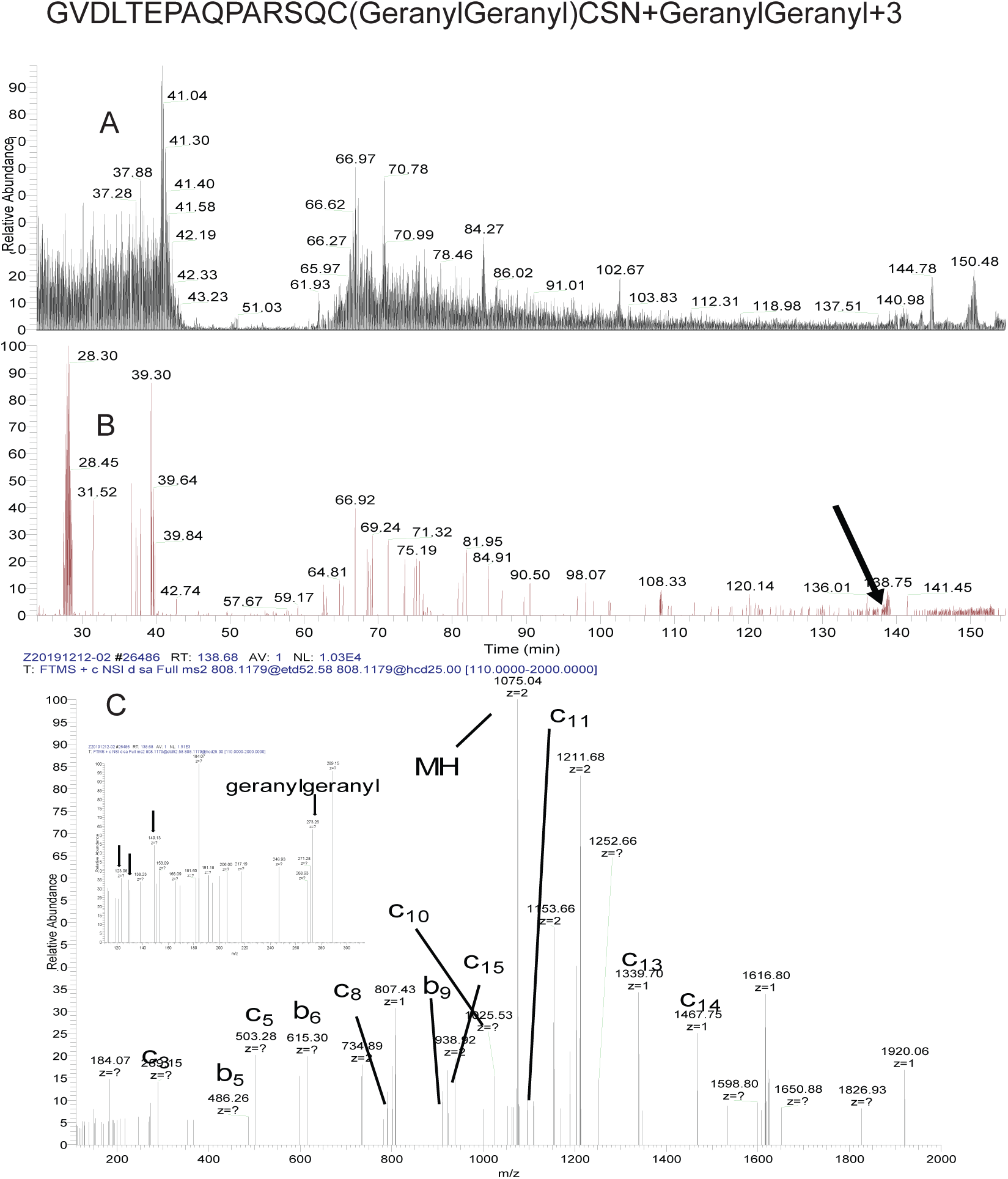
Elution of GVDLTEPAQPARSQCCSN (M+3H+ =807.45) from analytical C18 column. A. Base peak chromatogram from 155 min 20-60% acetonitrile gradient run. B. Extracted ion chromatogram showing (arrow) elution position of the peptide. C. ET-HCD MSMS spectrum showing b and c-type ions and low mass fragments (inset) derived from neutral loss of the geranylgeranyl moiety which also appears in an intact form in the spectrum at m/z 273.26.

## DISCUSSION

Despite their fundamental importance in cellular functions such as growth regulation, nuclear membrane structure/function, cytoskeletal rearrangement and cellular transport activities, global proteomic methods for characterization of prenylated proteins in unmodified cells and tissues have remained elusive. The current study describes a new approach designed to capture isoprene-modified peptides and proteins for further biological characterization of these important membrane-interacting sequences. The method takes advantage of novel chemistry associated with the thioether linked isoprene at the C-terminus of prenylated species and its interaction with 2-thiopyridyl-protected thiopropyl groups attached to Sepharose. The resin-attached group represents a relatively hydrophobic surface apart from the hydroxyl group. It has been demonstrated that disulfide groups are hydrophobic and, in proteins, are generally located within the hydrophobic core (23). This coupled with the hydrophobic nature of the thiopyridyl (24) group provides a potential mechanism for the prenyl bind-and-release mechanism uncovered in this study. The thiopropyl Sepharose resin used here is typically used to capture thiols through formation of disulfide bonds with the resin and was used by Forrester et al. to target palmitoylated peptides after chemical conversion of cysteine sulfur linked acyl palmitoylates to thiols in a method dubbed resin-assisted capture, “RAC” (25) which takes advantage of conventional bonding to the resin in contrast to the non-covalent interactions described here. Reduction of the disulfide in this study resulted in release of the thiopyridyl group and some isoprenylated peptides from the resin; further elution at low pH with 50% acetonitrile released more peptides. Results with a glutathione-modified thiopyridyl resin (glutathione is larger and more hydrophilic than the propyl group) were poor. Therefore, the chemistry at the resin surface including the presence of charged or polar groups undoubtedly plays a critical role in these phenomena. Although the exact mechanism of this interaction remains to be fully understood, further manipulation of the resin chemistry should provide more insight, perhaps even allowing enhanced performance.

Several interesting features emerged in the data (see Tables 1 and 2). Peptides from CN37, a major protein component of the myelin membrane and from CDC42, a regulator of spindle microtubule attachment during mitosis and of cell polarity and migration, were found in high abundance during the analysis. c-terminal peptides from these proteins were found in both farnesyl and geranylgeranyl forms. The presence of both modifications is consistent with a report that RhoB exhibits similar “flexibility” in its preference for the two isoprenes (26). The data presented here provides clear evidence that CN37 and CDC42 are modified by either FTase or GGTase I *in vivo.* We also found intermediate forms of the c-termini from CN37, CDC42 and guanine nucleotide binding proteins i.e. fully processed prenyl peptides with methyl ester modifications were found but also forms that were <u>only</u> prenylated. This is the first report to our knowledge of mixed prenylation of CDC42. The retention times of peptides on the C18 column demonstrated the difference in polarity conferred by the presence of a geranylgeranyl group versus a farnesyl (figure 4). The meaning of the presence of both isoprenes on the same protein is still unclear at this point although others (27) have speculated that the presence of farnesyl or geranylgeranyl may confer some specificity to membrane association and it was reported that both CN37 isoforms could be labelled by either ^3^H farnesyl pyrophosphate or ^3^H geranylgeranyl pyrophosphate *in vitro* or in C6 cells after treatment with HMG Co A reductase inhibitors and labelling as above (28). RhoB was shown to be modified by both types of prenyl groups using a cell line that overexpressed the recombinant RhoB protein and labelling with ^3^H mevalonate (29). Subsequent reports (30, 31) detailed a “gain of function” for Rho-B after inhibition of FT (and subsequent geranylgeranylation of the molecule) in Rat1/ras transformed cells such that the geranylgeranylated molecule drove reversion of the transformed phenotype and inhibition of the cell cycle. Although the possible functional biological significance of mixed prenylation of CN37 and CDC42 is unknown at present, enrichment and analysis by LCMS demonstrated in the current study opens the door to direct analysis of these phenomena in tissue samples. In addition to the mixed prenyl forms detected above, intermediate forms of targeted c-terminal peptides were detected. Isoprene-modified peptides were found with the “AAX” sequence removed but not methylated. In contrast with the recent report by Storck et al. (19) we only found a single example of a CAAX box c-terminal peptide (which appeared in the peptides released from the resin by chymotryptic digestion alone; data not shown) with an intact CAAX sequence; otherwise, the “AAX” sequence had always been removed by the Rce I enzyme. This difference may result from kinetic differences in prenyl protein processing between labelled samples in culture versus the more “steady state” represented by the tissue samples from mouse brain or simply differences in our capture techniques. Surprisingly, c-terminal peptides with a methyl group present on the c-terminal cysteine and no prenylation were reported in the database search results. However, we believe that these were formed in each case from a neutral loss of the isoprene group in the source of the mass spectrometer based on their detection only when a prenylated equivalent was eluting. We observed this phenomenon primarily in the highly abundant peptides, suggesting that the neutral loss may not represent a major problem in prenylated peptide analysis using mass spectrometry. However, altering instrument parameters to reduce the phenomenon, such as by lowering the temperature of the entry capillary into the instrument may be advisable, although this has not yet been tested.

We detected c-terminal peptides from proteins recognized by the CAAX protein enzymes FTase and GGTase I using HCD and in longer runs using EThcD c-terminal peptides from Rab-3a, Rab3c, Rab7a, Rab14 and Rab5a that were modified by 2 geranylgeranyl isoprenes (catalyzed by GGTase2) were identified. Many peptides from other Rab proteins were seen (Table S1) suggesting that detection of these peptides may also be achieved using higher starting acetonitrile in gradient elutions (20 vs 10 percent) coupled with enhanced fragmentation capability offered by EThcD for large peptides. Due to their size and hydrophobicity, the peptides from doubly geranylgeranylated c-termini migrated extremely late from the C18 column, even with the latter chromatographic adjustments, suggesting that alternate chromatographic approaches (e.g. C4 or other more polar stationary phases) could help in separation and characterization of these peptides. Further experiments will help to define optimal combinations of the above.

In summary, we developed a method that is capable of capturing prenylated proteins directly from tissue samples and that using high-resolution LCMS coupled with customized informatics, we were able to identify and characterize tandem mass spectra from native c-terminal peptides derived from a range of these molecules for the first time. Guided by the current experiments, a combination of enhanced resin design with optimized chromatographic and mass spectrometric approaches should enhance the range of peptides identified and characterized. This raises for the first time the possibility that changes in the prenylome of clinical samples or other non-cultured cells and tissues can be directly assessed with minimal pre-processing. Another important area of interest will be tissue responses to pathogen infection which is known in some cases to involve key participation of the prenylation pathway of host cells (e.g. (32).

## Acknowledgements

Financial support was provided by the Dr Mariam and Sheldon Adelson Medical Research Foundation (to ALB). We further acknowledge a UCSF School of Pharmacy, 2019 Mary Anne Koda-Kimble Seed Award for Innovation. The authors would also like to acknowledge the generous contribution of prenylated A-Factor peptides by Professor Fred Nader at the College of Staten Island, City University of New York. These peptides were critical to our understanding of the effect of prenylation on peptide behavior. We further acknowledge Bob Nairn of Nouryon Pulp & Performance Chemicals for generously supplying all of the Kromasil C18 and C4 resins used in these studies. We also thank Juan A. Oses-Prieto for many valuable technical discussions.

## Data Availability

Data are available using Protein Prospector through the MS Viewer application (http://msviewer.ucsf.edu/prospector/cgi-bin/msform.cgi?form=msviewer) using the search keys lskwuid29u and hyzecfvqkp.

